# The landscape of metabolic brain alterations in Alzheimer’s disease

**DOI:** 10.1101/2021.11.15.468698

**Authors:** Richa Batra, Matthias Arnold, Maria A. Wörheide, Mariet Allen, Xue Wang, Colette Blach, Allan I. Levey, Nicholas T. Seyfried, Nilüfer Ertekin-Taner, David A. Bennett, Gabi Kastenmüller, Rima F. Kaddurah-Daouk, Jan Krumsiek, Alzheimer’s Disease Metabolomics Consortium (ADMC)

## Abstract

**INTRODUCTION:** Alzheimer’s disease (AD) is accompanied by metabolic alterations both in the periphery and the central nervous system. However, so far, a global view of AD-associated metabolic changes in brain has been missing.

**METHODS:** We metabolically profiled 500 samples from the dorsolateral prefrontal cortex. Metabolite levels were correlated with eight clinical parameters, covering both late-life cognitive performance and AD neuropathology measures.

**RESULTS:** We observed widespread metabolic dysregulation associated with AD, spanning 298 metabolites from various AD-relevant pathways. These included alterations to bioenergetics, cholesterol metabolism, neuroinflammation and metabolic consequences of neurotransmitter ratio imbalances. Our findings further suggest impaired osmoregulation as a potential pathomechanism in AD. Finally, inspecting the interplay of proteinopathies provided evidence that metabolic associations were largely driven by tau pathology rather than β-amyloid pathology.

**DISCUSSION:** This work provides a comprehensive reference map of metabolic brain changes in AD which lays the foundation for future mechanistic follow-up studies.

## 1 Background

Alzheimer’s disease (AD) is the most common cause of dementia, with prevalence rates expected to increase markedly over the next decades (1). AD is a neurodegenerative disorder defined by the deposition of β-amyloid and accumulation of neurofibrillary tangles of phosphorylated tau protein in the brain (2). These proteinopathies are accompanied by other pathogenic processes, including neuroinflammation, oxidative stress, innate immune response, and neurotransmission (1). A large body of evidence further implicates metabolic pathways both in the periphery and in the central nervous system in AD (3–8). Notably, metabolic enzymes and transporters are among the most commonly targeted proteins in pharmaceutical interventions across all diseases (9, 10), emphasizing the translational potential of systematically identifying metabolic alterations. However, until now a comprehensive reference map of metabolic brain changes related to AD, AD-associated neuropathological manifestation, and cognitive decline has been missing.

To fill this gap, we here present a large, multi-center study from the Accelerating Medicine Partnership in AD (AMP-AD) consortium, analyzing a total of 500 post-mortem brain tissue samples from the dorsolateral prefrontal cortex (DLPFC) using broad, non-targeted metabolomics measurements. This dataset represents, to the best of our knowledge, the largest metabolomics study of aging brain tissue to date. In the present paper, we first provide an overview of the extensive metabolic changes in brain, spanning multiple AD-related traits, including neuropathological β-amyloid and tau tangle burden, as well as late-life cognitive performance. This is followed by replication analysis in independent samples from different brain regions.

We then present several examples of how such a large dataset can be used to extract novel metabolic insights into AD from the vast number of identified associations. These include: (1) Extending metabolic characterization of pathways that previously have been implicated in AD, including bioenergetic pathways, cholesterol metabolism, and neuroinflammation. (2) Establishing broad impairment of osmoregulation as a potentially relevant pathomechanism in AD. (3) Endorsing the concept of an imbalance between excitatory/inhibitory neurotransmitter ratios in AD by integrating our metabolomic findings with additional proteomic data. (4) Identifying tau loadas a potential driver of metabolic dysfunction in the AD brain, with minimal contributions from β-amyloid load.

Finally, to maximize utilization of our study by the scientific community, we have made our data, code, and findings available through the AD Knowledge Portal and an interactive web resource at https://omicscience.org/apps/brainmwas/.

## 2 Methods

### 2.1 Cohorts, clinical data, and neuropathological data

#### 2.1.1 ROS/MAP cohort

The Religious Order Study (ROS) and Rush Memory and Aging Project (MAP) cohorts (11, 12) are two longitudinal, clinicopathologic studies conducted by the Rush Alzheimer’s Disease Center. ROS started in 1994 with the recruitment of individuals from religious communities across the United States. MAP started in 1997 with the recruitment of individuals from a wide range of backgrounds and socio-economic statuses from northeastern Illinois. Both cohorts were approved by an institutional review board of Rush University Medical Center. Both studies focus on older individuals who agreed to longitudinal clinical analysis and brain donation after death. All participants signed an informed consent, an Anatomic Gift Act, and a repository consent to allow their data and biospecimens to be shared. Following enrollment in the study, participants were evaluated for physical and cognitive function annually. After death, pathologic assessment was performed. Initially, 514 samples from the DLPFC brain region were used for metabolomics profiling, along with associated metadata, including medications taken during lifetime, age at death, sex, BMI, postmortem interval, *APOE* genotype status, education history, cognitive scores during lifetime, cognitive decline (computed-based on longitudinal cognitive scores), clinical diagnosis at death, β-amyloid and paired helical filament (PHF)-tau protein load in brain tissue, global burden of AD neuropathology (mean of neuritic plaques, diffuse plaques, and neurofibrillary tangles), NIA-Reagan score, Braak stage, and CERAD score. Neuropathological diagnosis was derived using the following criteria: AD case status was assigned where Braak stage was ≥ 4 and CERAD score was ≤ 2; control case status was assigned where Braak stage was ≤ 3 and CERAD score was ≥ 3. All clinical parameters have previously been described in detail (13). All 514 participants were of Caucasian descent. Of note, the *APOE*-ε4 allele frequency in the overall ROS/MAP cohort is 13.27%, as determined from the ROS/MAP online data repository for 2,097 non-Hispanic white participants (14). This number is in line with other reported ε4 allele frequencies in Caucasian populations (15–17). Since our metabolomics sub-cohort of 514 samples was enriched for AD cases (44%), the ε4 frequency in our study with 25% was subsequently higher than the background.

#### 2.1.2 Mayo Clinic cohort

Initially, and before filtering, 84 samples from the temporal cortex brain region were obtained from the Mayo Clinic Brain Bank. Details on this cohort have been provided in previous studies (18, 19). All samples received diagnoses at autopsy following neuropathologic evaluation. 64 samples had a neuropathologic diagnosis of AD with Braak :=4.0 and 20 control samples had Braak :′,3 and without any neurodegenerative diagnoses. All 84 samples were from North Americans of European descent with ages at death :=60 for AD and :=53 for controls.

#### 2.1.3 Cohort differences

Three cohorts were used in this publication – ROS/MAP for discovery, Mayo brain clinical cohort for replication, and a published Baltimore Longitudinal Study of Aging (BLSA) based study (5) for comparison. These cohorts have fundamental differences: (a) Participant recruitment. BLSA is an aging study, Mayo Clinic samples are from an archival brain bank with neuropathologic diagnoses of AD and control, while ROS/MAP recruited older people. (b) AD-related traits. Mayo has diagnosis determined by neuropathology and BLSA has diagnosis determined based on neuropathology and cognitive conditions. ROS/MAP records several neuropathological as well as cognitive scores. (c) Unlike the other two cohorts, ROS/MAP collects various lifetime variables longitudinally, including cognitive scores, lifestyle, medications taken by participants. (d) Sample sizes were lower in BLSA (n = 43), and Mayo (n = 84), compared to ROS/MAP (n = 514). (e) Different brain regions were profiled. BLSA sampled frontal and temporal gyrus, Mayo the superior temporal gyrus of the temporal cortex, and ROS/MAP the dorsolateral prefrontal cortex (DLPFC).

### 2.2 Metabolomics profiling

Brain metabolic profiles were measured using the untargeted metabolomics platform from Metabolon Inc. Briefly, tissue samples were divided into four fractions; two for ultra-high performance liquid chromatography-tandem mass spectrometry (UPLC-MS/MS; positive ionization), one for UPLC-MS/MS (negative ionization), and one for a UPLC-MS/MS polar platform (negative ionization). The combination of these four runs ensures a broad coverage of hydrophilic and hydrophobic substances. Peaks were quantified using the area under the curve in the spectra. To account for run-day variations, peak abundances were normalized by their respective run-day medians. Compounds were identified using an internal spectral database. A detailed description of all experimental procedures can be found in the **supplementary information.**

### 2.3 Data preprocessing

#### 2.3.1 ROS/MAP and Mayo metabolomics

Metabolites with over 25% missing values were filtered out, leaving 667 out of an original 1,055 metabolites for ROS/MAP and 664 out of 827 for Mayo. Probabilistic quotient normalization was applied to correct for sample-wise variation (20), followed by log_2_ transformation. Remaining missing values were imputed using a k-nearest-neighbor-based algorithm (21). Outlier samples in the data were removed using the local outlier factor method (22) implemented in the R package bigutilsr. To account for remaining irregularly high or low single concentrations, values with absolute abundance above *q = abs(qnorm(0.0125/n*)), with *n* representing the number of samples, were set to missing. This formula finds the cutoff for values with less than 2.5% two-tailed probability to originate from the same normal distribution as the rest of the measurement values, after applying a Bonferroni-inspired correction factor (division by sample size). These new missing values were then imputed by another round of the k-nearest-neighbor algorithm.

#### 2.3.2 ROS/MAP proteomics

Proteomics data was downloaded from the AMP-AD Knowledge Portal (https://adknowledgeportal.synapse.org), details of proteomic profiling and data processing can be found in the original publication (23). Briefly, data were log_2_-transformed and corrected for batch effects using the ‘median polish’ approach. In our analysis, proteins with over 25% missing values were filtered out, and remaining missing values were imputed using a k-nearest-neighbor-based algorithm (21). Outliers were treated with the same approach as the metabolomics data (see above).

#### 2.3.3 Medication correction

For the ROS/MAP cohort, all prescription and over-the-counter medications were collected at each study visit. To account for influences of these medications on metabolomics and proteomics, a linear stepwise backward selection approach was used (4). All preprocessing steps were performed using the maplet R package (24).

### 2.4 Differential analysis of metabolites and proteins

#### 2.4.1 ROS/MAP

Five outliers identified by the local outlier factor method, six samples with missing medication information, 1 sample with missing BMI, and 2 samples with missing *APOE* genotype status were removed from further analysis. Therefore, after preprocessing, 500 samples were used for metabolic analysis, and 262 matching samples were used for proteomic analysis. Metabolite and protein associations were computed using generalized linear models with the traits as response variables and molecule levels as predictors. For statistical analysis, the following transformations were made: Square root of β-amyloid load and binarized NIA-Reagan score (0 – low likelihood of AD, 1 – high likelihood of AD). For the association analysis with clinical diagnosis, 8 non-AD-related dementia samples were removed. The trait ‘cognitive decline’ is based on the slope of cognitive values over time. In order for higher cognitive decline to be represented by higher values, the direction of this slope was inverted during the analysis. Appropriate link functions were used according to the respective variable types, i.e., identity link function for continuous traits (regular linear regression for β-amyloid, tau tangles, global burden of pathology, cognition levels, cognitive decline), logit for binary traits (logistic regression for NIA-Reagan score and NP diagnosis), and probit for the ordinal trait (ordinal regression for clinical diagnosis after death). All models accounted for confounding effects of age, sex, BMI, postmortem interval, number of years of education, and number of *APOE*-ε4 alleles. Notably, age, sex, years of education did not show much influence on metabolic profiles of cognitively normal samples but are known confounders of AD (3), justifying the correction in the models. To account for multiple hypothesis testing, p-values were corrected using the Benjamini-Hochberg (BH) method (25). Cognitive levels are inversely related to AD, and thus the direction of associations with the variable ‘cognition’ was reversed after statistical analysis.

#### 2.4.2 Mayo

One AD sample with missing *APOE* genotype status was removed from further analysis. Therefore, after preprocessing, 63 AD and 20 control samples with complete information on age at death, *APOE-*ε4 allele status, and sex were used for our analysis. For replication, metabolites that were associated with any of the eight AD-related traits in the ROS/MAP cohort at 5% FDR were selected. The analysis was performed using two subsequent logistic regressions with diagnosis as outcome. The first model was built without any confounder correction. To account for multiple hypothesis testing, p-values were corrected using the Benjamini-Hochberg (BH) method (25). Metabolites with adjusted p-values < 0.05 were selected for the second model. The second model was built with confounders sex, number of *APOE*-ε4 alleles, and age at death. Metabolites with nominal p-values < 0.05 in the second model were considered replicated. All analyses were performed using the maplet R package (24).

### 2.5 Stratified analysis

To determine the influence of sex and *APOE*-ε4 status on metabolic associations, we performed a stratified analysis per factor (sex and *APOE*-ε4 status) for each AD-related trait. Metabolites significant at 5% FDR were selected to compute within-group (male/female, *APOE*-ε4 positive/ *APOE*-ε4 negative) metabolic associations with AD-related traits. 𝛽 estimates across groups per metabolite were compared using z-scores (26), defined as 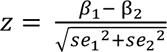 where 𝛽 and 𝛽 are the 𝛽 coefficients from the linear regressions performed in the two groups, and 𝑠𝑒_)_ and 𝑠𝑒_*_ are the corresponding standard errors. Z-scores are approximately standard normally distributed and were thus used to compute p-values using a normal distribution. Any metabolite with a nominal p-value < 0.05 was considered significantly different within the respective group.

### 2.6 Metabolic network inference

To infer the metabolite-metabolite interaction network, a partial correlation-based Gaussian graphical model (GGM) was computed using the GeneNet R package (27). P-values of partial correlations were corrected using the Bonferroni method. Partial correlations with adjusted p-values < 0.05 were used for network construction between metabolites. To annotate the metabolic network with AD associations, a score was computed for each metabolite/trait combination as follows: 𝑝*_score_* = 𝑑 ∗ (−1 ∗ log_10_(𝑝. 𝑎𝑑𝑗)), where 𝑝. 𝑎𝑑𝑗 is the adjusted p-value of the model, and *d* is the direction (−1/1) of metabolite association based on the test statistic (positive or negative correlation with AD-related trait). To aggregate the signal across the traits, an overall score was defined as the *p_score_* with maximum absolute value. This overall score was used to color the nodes in GGM in Figure 2f and the online supplement.

## 3 Results

### 3.1 Cohort description and characteristics of brain metabolomics data

We analyzed brain samples of 500 participants from the Religious Order Study and the Rush Memory and Aging Project (ROS/MAP) cohorts(11, 12), including 352 females and 148 males, with a mean age at death of 91 (**Table 1**). Following enrollment in the study, participants were evaluated for physiological and cognitive function once per year (**Figure 1**). Neuropathology was assessed after autopsy. Out of the 500 participants, 220 were diagnosed with AD (with or without a secondary cause of dementia) at the time of death, 119 had mild cognitive impairment, 153 were without cognitive impairment, and 8 had other forms of dementia. Samples from the dorsolateral prefrontal cortex (DLPFC) brain region were used for untargeted metabolic profiling. Metabolomics measurements were analyzed in relation to eight AD-related traits covering late-life cognitive assessments and postmortem pathology: Clinical diagnosis at time of death, level of cognition proximate to death, cognitive decline during lifetime, β-amyloid load, tau tangle load, global burden of AD pathology (global NP), NIA-Reagan score, neuropathological diagnosis inferred based on a combination of Braak stage and CERAD scores (NP diagnosis; see methods for diagnostic criteria). A detailed description of these eight AD-related traits is provided in Table 2.

**Figure 1:**
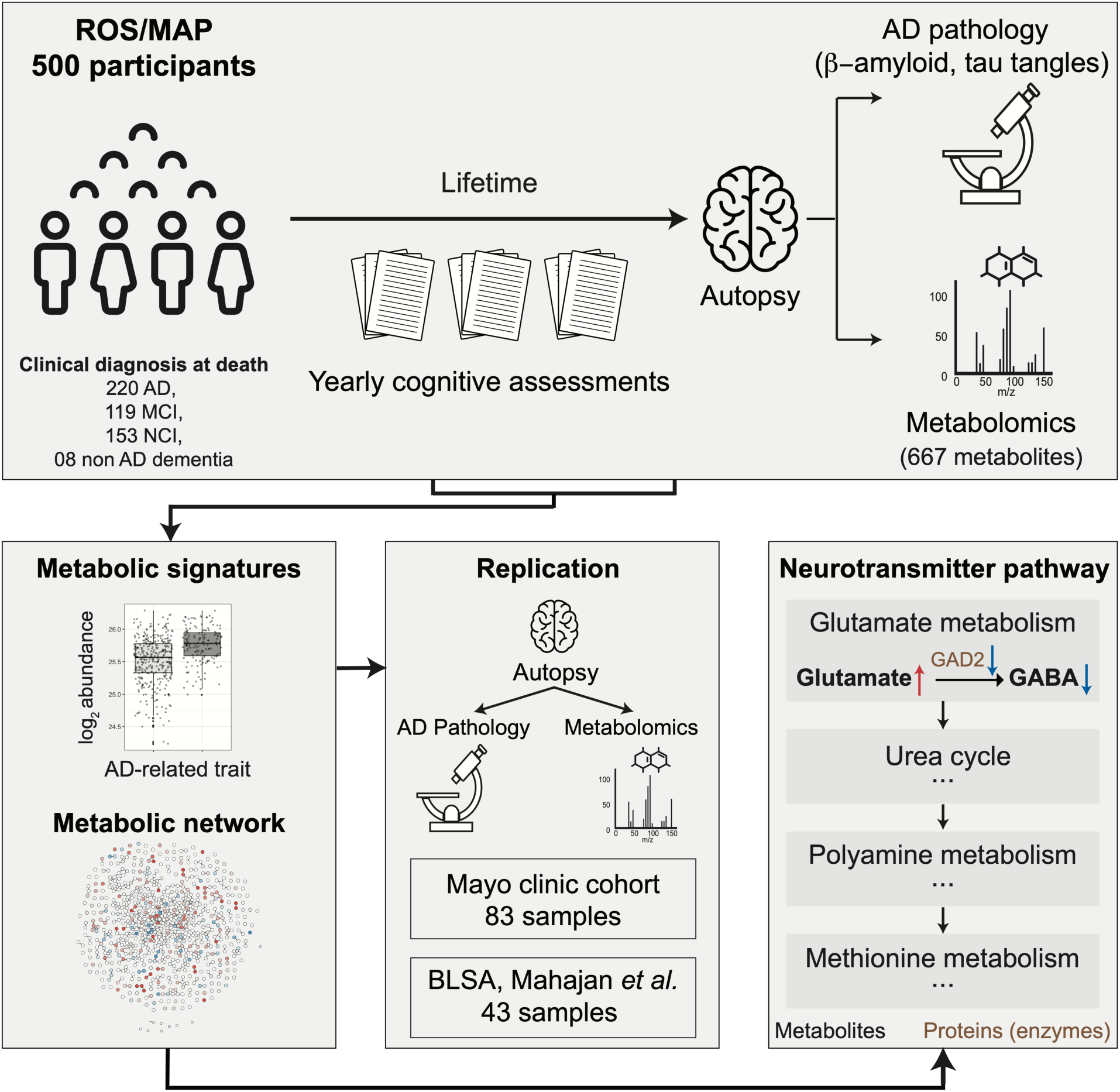
Study overview. 500 ROS/MAP participants were included in this analysis. For each participant, data was available on cognitive assessments during lifetime, postmortem AD brain pathology, and brain metabolic profiles from the dorsolateral prefrontal cortex (DLPFC) region. Metabolic profiles were investigated for associations with AD-related traits and a metabolite interaction network was inferred using a Gaussian graphical model (GGM). Associations were tested for replication in 83 temporal cortex samples from the Mayo Clinic brain bank cohort and compared to a previously published brain-based study. Finally, various pathways previously implicated in AD were metabolically characterized, and a detailed metabolomic/proteomic characterization of the glutamate/GABA neurotransmitter pathway was generated.

**Table 1.** ROS/MAP cohort overview. Postmortem interval refers to the time between death and sample preservation. BMI = body mass index. IQR = interquartile range, i.e., middle 50% of the data.

|  |  |
| --- | --- |
| **Total samples** | **N = 500** |
| **Sex** |  |
| Female | 352 (70.4%) |
| Male | 148 (29.6%) |
| **Postmortem interval (hours)** | 6.6 (5.2, 8.7) |
| **BMI** | 25.2 (22.5, 28.2) |
| **Years of education** | 16.00 (13.00, 18.00) |
| **Age at death** | 91 (87, 95) |
| **APOE4 alleles** |  |
| 0 | 374 (75%) |
| 1 | 121 (24%) |
| 2 | 5 (1%) |
| **Clinical diagnosis at death** |  |
| AD | 220 (44%) |
| Mild cognitive impairment (MCI) | 119 (23.8%) |
| No cognitive impairment (NCI) | 153 (30.6%) |
| Other | 8 (1.6%) |
| Format: N (%) or median (IQR) |  |

**Table 2:** Description of these AD-related traits investigated in this study.

| **AD-related trait** | **Description of trait** | **Trait data type** |
| --- | --- | --- |
| β-amyloid | Immunohistochemistry-based overall β-amyloid load | numeric |
| Tau tangles | Immunohistochemistry-based overall paired helical filament (PHF)-tau tangles load | numeric |
| Global NP | Summary of pathology derived from counts of: neuritic plaques, diffuse plaques, and neurofibrillary tangles | numeric |
| NIA Reagan | NIA Reagan diagnosis of Alzheimer's disease derived from Braak and CERAD scores | binary |
| NP Diagnosis | Mayo clinic diagnosis of Alzheimer's disease derived from Braak and CERAD scores | binary |
| Clinical Diagnosis | Consensus cognitive diagnosis at time of death | ordinal |
| Cognition | Global cognitive function determined at the last timepoint before death | numeric |
| Cognitive Decline | Rate of change in global cognition over time | numeric |

The metabolomics platform identified 667 metabolites from various chemical classes (super-pathways) in the brain samples, including lipids (42.7%), amino acids (22.6%), nucleotides (6.7%), carbohydrates (6.3%), cofactor and vitamins (4.3%), xenobiotics (3.7%), peptides (2.1%), and energy-related metabolites (1.5%) **(Figure 2a, Supplementary Table 1)**. Previous blood-based metabolomics studies reported strong influences of medications and supplements (such as vitamins) on metabolic profiles (4). To investigate such effects in brain tissue-based metabolic profiles, we examined influences of 103 grouped medication classes and supplements on metabolic abundances. 552 out of 667 (82.75%) of the metabolites correlated with one or more medications or supplements taken during lifetime. The group of medications to treat benign prostatic hypertrophy associated with the highest number of metabolites (81 metabolites), followed by diuretics (55 metabolites) and multivitamins (52 metabolites). A comprehensive list of medication classes and their effect on the metabolome is provided in **Supplementary Table 2**. Given these strong associations with the metabolome, medication effects excluding AD-related and neurologic drugs were regressed out from the metabolic profiles for all following analyses. Moreover, since the postmortem interval (PMI) before sample collection at autopsy may also impact analyte levels, we investigated its effects and found that 307 metabolites associated with PMI (**Supplementary Table 3**). PMI was therefore included as a covariate in all following analyses.

**Figure 2:**
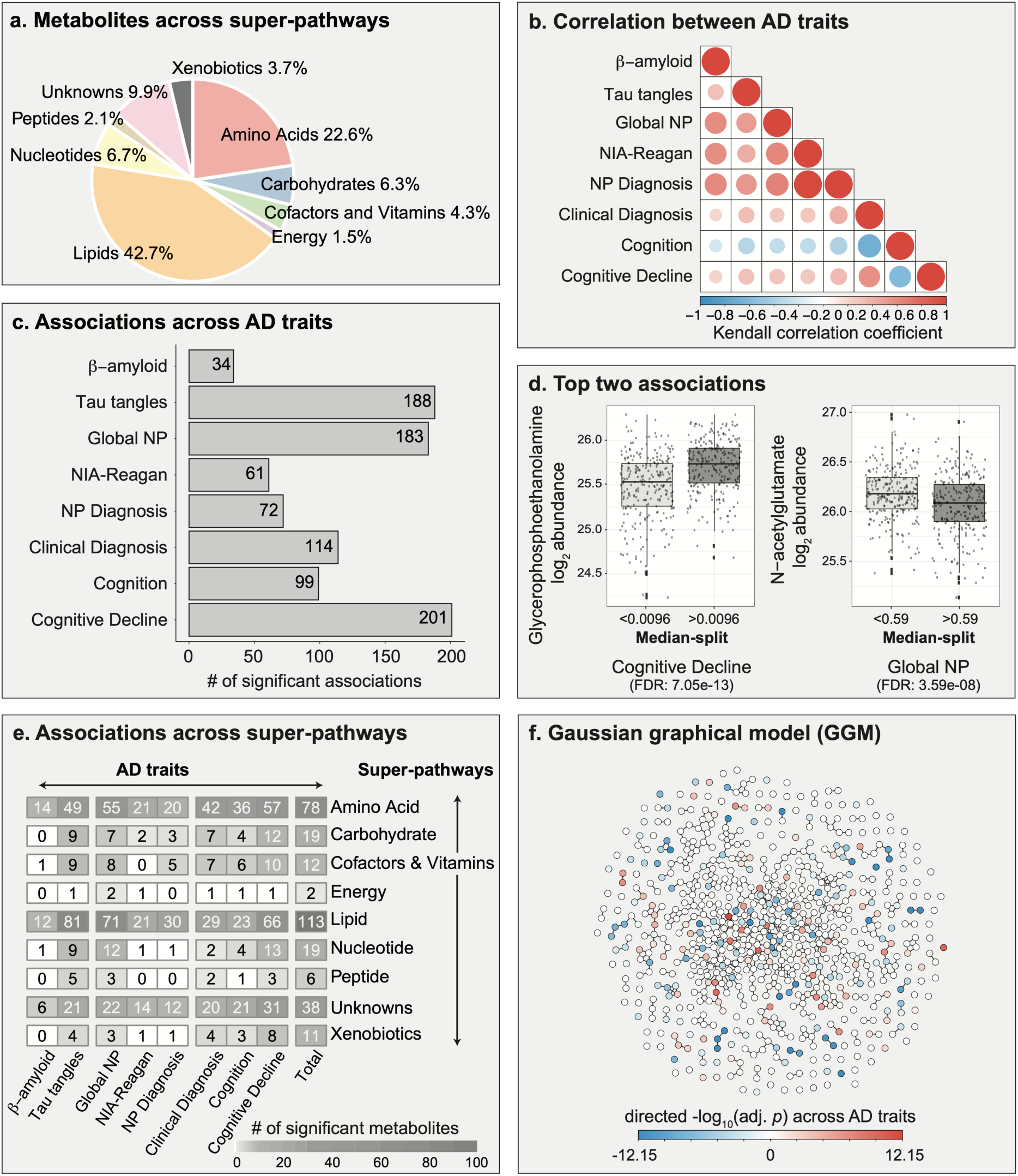
Overview of metabolic associations with AD traits. **a**, Metabolites measured in brain samples are distributed across various metabolic classes, referred to as “super-pathways” throughout the manuscript. **b**, Kendall correlations across the eight AD-related traits. **c**, A total of 298 metabolites were associated with at least one of the eight AD-related traits. **d**, Examples of two metabolites with the lowest adjusted p-values. Note that the traits were discretized (median split) for visualization. **e**, Distribution of metabolic associations across super-pathways. **f**, Gaussian graphical model of metabolites. Metabolites are colored based on the negative log_10_ of the lowest adjusted p-value across AD-related traits multiplied with the direction of the respective effect estimate. Global NP = Global burden of AD neuropathology. NP Diagnosis = postmortem diagnosis based on Braak stage and CERAD score. Cognition = cognitive assessment scores from last premortem test. Cognitive Decline = Change in cognitive assessment scores over time until death.

To obtain a baseline understanding of metabolism in brain, we used cognitively normal samples to compute associations between measured metabolites and demographic parameters *independent of AD pathology*. This included metabolic associations with age, BMI, sex, and years of education (as a proxy for socioeconomic status). Two metabolites, 1-methyl-5-imidazoleacetate and N6-carboxymethyllysine, were significantly associated with age at 5% false discovery rate (FDR). Surprisingly, there were no significant associations with sex, BMI, or education, which is in stark contrast to findings in blood (28–30). Details of this baseline analysis can be found in **Supplementary Table 4.**

### 3.2 AD is associated with widespread metabolic alterations in brain

To assess AD-related metabolic changes, we computed statistical associations between metabolic profiles and the 8 AD-related traits. All statistical models accounted for AD-related confounders (age, sex, years of education, BMI, and copies of APOE4) as well as postmortem interval. A total of 298 out of 667 metabolites (44.7%) were significantly associated with one or more AD traits at 5% false discovery rate (FDR). 80 out of the 298 metabolites showed unique associations with just one of the traits. A total of 218 metabolites were associated with more than one trait, which is likely due to high correlations across traits (**Figure 2b, Supplementary Figure 1**). The majority of the 298 metabolites was associated with one of three AD traits: Cognitive decline (n = 201), tau tangles (n = 188), and global burden of pathology (n = 183) (**Figure 2c**). Interestingly, only 34 metabolites associated with β-amyloid, which was the lowest number of associations among the eight AD traits. Furthermore, we observed that 159 out of the 298 metabolites (53.4%) were associated with both premortem parameters and postmortem pathological assessments. All statistical results are provided in **Supplementary Table 5**. Sex-based stratified analysis revealed that 29 of the 298 metabolites (10%) showed associations with at least one trait that were significantly modulated by sex (**Supplementary Table 6**), and APOE4-stratified analysis showed that associations of 77 metabolites (26%) were influenced by APOE4 status (**Supplementary Table 7**).

The three most significantly changed metabolites for each of the eight AD-related traits are listed in **Table 3**. Moreover, we illustrate the two most significant associations in the dataset as visual examples: Glycerophosphoethanolamine (GPE) levels, which positively associated with cognitive decline (FDR: 7.05e-13, **Figure 2d, left**), and N-acetylglutamate, which negatively associated with global AD pathology (FDR: 3.59e-08, **Figure 2d**, **right**). GPE levels were higher with lower cognitive abilities, which corroborates previously published findings(31). N-acetylglutamate levels showed lower levels with higher AD pathology load.

**Table 3.**
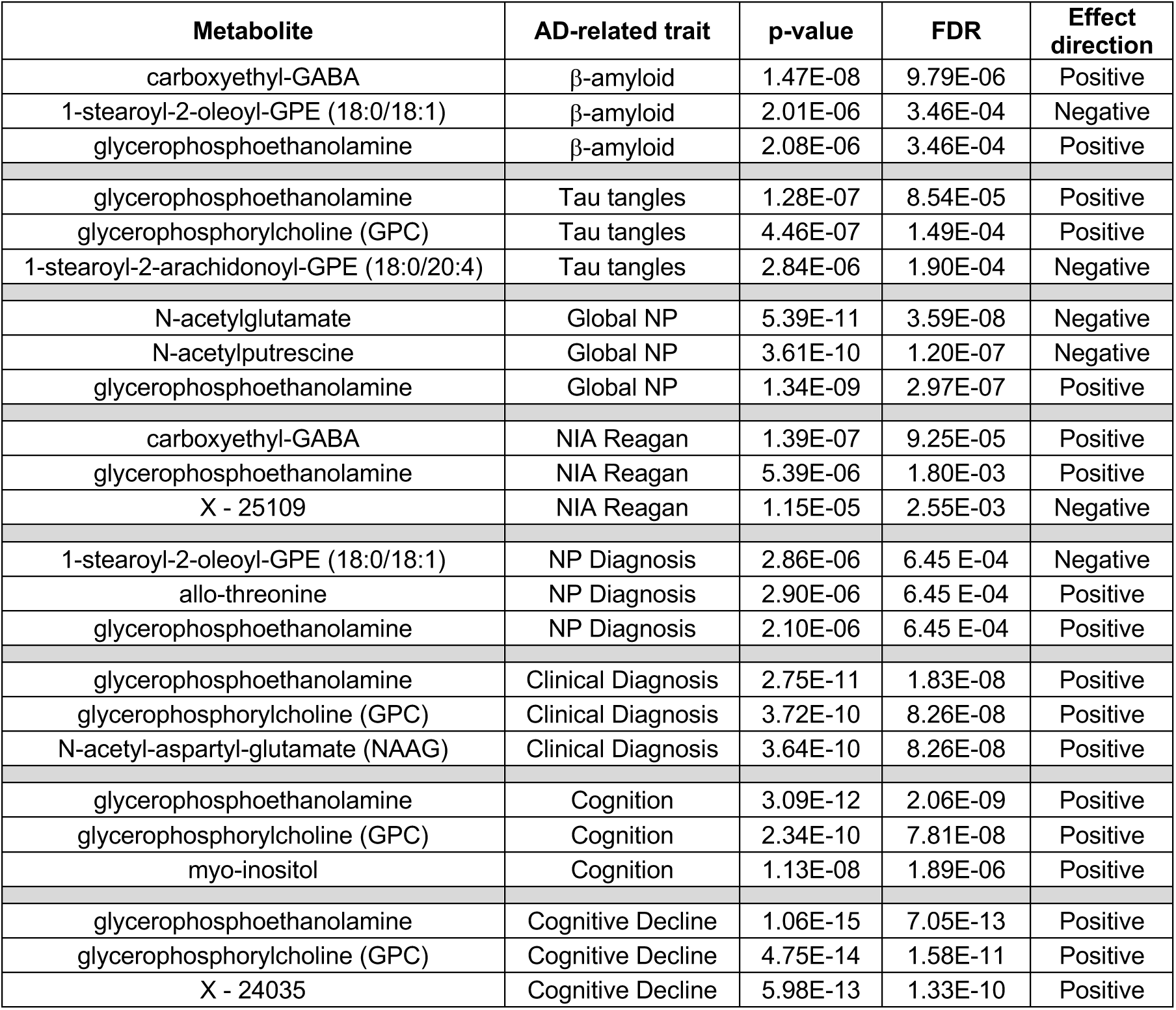
Top three metabolites associated with each AD-related trait. Global NP = Global burden of AD neuropathology. NP Diagnosis = postmortem diagnosis based on Braak stage and CERAD score. Cognition = cognitive assessment scores from last premortem test. Cognitive Decline = Change in cognitive assessment scores over time until death.

| Metabolite | AD-related trait | p-value | FDR | Effect direction |
| --- | --- | --- | --- | --- |
| carboxyethyl-GABA | β-amyloid | 1.47E-08 | 9.79E-06 | Positive |
| 1-stearoyl-2-oleoyl-GPE (18:0/18:1) | β-amyloid | 2.01E-06 | 3.46E-04 | Negative |
| glycerophosphoethanolamine | β-amyloid | 2.08E-06 | 3.46E-04 | Positive |
| glycerophosphoethanolamine | Tau tangles | 1.28E-07 | 8.54E-05 | Positive |
| glycerophosphorylcholine (GPC) | Tau tangles | 4.46E-07 | 1.49E-04 | Positive |
| 1-stearoyl-2-arachidonoyl-GPE (18:0/20:4) | Tau tangles | 2.84E-06 | 1.90E-04 | Negative |
| N-acetylglutamate | Global NP | 5.39E-11 | 3.59E-08 | Negative |
| N-acetylputrescine | Global NP | 3.61E-10 | 1.20E-07 | Negative |
| glycerophosphoethanolamine | Global NP | 1.34E-09 | 2.97E-07 | Positive |
| carboxyethyl-GABA | NIA Reagan | 1.39E-07 | 9.25E-05 | Positive |
| glycerophosphoethanolamine | NIA Reagan | 5.39E-06 | 1.80E-03 | Positive |
| X - 25109 | NIA Reagan | 1.15E-05 | 2.55E-03 | Negative |
| 1-stearoyl-2-oleoyl-GPE (18:0/18:1) | NP Diagnosis | 2.86E-06 | 6.45 E-04 | Negative |
| allo-threonine | NP Diagnosis | 2.90E-06 | 6.45 E-04 | Positive |
| glycerophosphoethanolamine | NP Diagnosis | 2.10E-06 | 6.45 E-04 | Positive |
| glycerophosphoethanolamine | Clinical Diagnosis | 2.75E-11 | 1.83E-08 | Positive |
| glycerophosphorylcholine (GPC) | Clinical Diagnosis | 3.72E-10 | 8.26E-08 | Positive |
| N-acetyl-aspartyl-glutamate (NAAG) | Clinical Diagnosis | 3.64E-10 | 8.26E-08 | Positive |
| glycerophosphoethanolamine | Cognition | 3.09E-12 | 2.06E-09 | Positive |
| glycerophosphorylcholine (GPC) | Cognition | 2.34E-10 | 7.81E-08 | Positive |
| myo-inositol | Cognition | 1.13E-08 | 1.89E-06 | Positive |
| glycerophosphoethanolamine | Cognitive Decline | 1.06E-15 | 7.05E-13 | Positive |
| glycerophosphorylcholine (GPC) | Cognitive Decline | 4.75E-14 | 1.58E-11 | Positive |
| X - 24035 | Cognitive Decline | 5.98E-13 | 1.33E-10 | Positive |

The 298 metabolites that were associated with AD-related traits were distributed across all super-pathways, including 113 (37.92%) within the largest super-pathway of lipids, followed by amino acids with 78 (26.17%) associations, and the rest in the remaining six super-pathways (**Figure 2e**). At the more fine-grained sub-pathway level, metabolic associations were distributed across 72 out of the 101 sub-pathways covered in the data (**Supplementary Figure 2**).

We statistically inferred a metabolic network and annotated it with effect directions and the lowest adjusted p-value across the eight AD traits (**Figure 2f**). The network is based on a Gaussian graphical model (GGM), which corresponds to a data-driven representation of biochemical pathways (32, 33). GGMs have previously been used to systematically investigate various trait effects on the metabolome (21, 28). To further explore our findings networks for each AD trait are available as a Cytoscape file (**Supplementary File 1**), as well as an interactive online version at https://omicscience.org/apps/brainmwas/.

Taken together, this analysis revealed global metabolic changes with respect to various AD-related clinical and neuropathological traits. These alterations encompass all measured metabolic super-pathways, highlighting the massive impact of the disease on brain metabolism.

### 3.3 AD-associated metabolic alterations overlap across independent brain studies

To strengthen confidence in our findings, we performed replication analysis using a metabolomics dataset from a Mayo Clinic brain bank cohort. In addition, we compared our results to a previously published brain metabolomics study based on samples from the Baltimore Longitudinal Study of Aging (BLSA) (5). Detailed replication results of Mayo can be found in **Supplementary Table 8** and published BLSA results used for comparison can be found in **Supplementary Table 9**.

In the Mayo data, 83 temporal cortex brain samples were used for untargeted metabolic profiling, including 63 AD patients and 20 controls. Of the 8 AD-related traits used in the discovery phase with ROS/MAP cohort, neuropathology-based diagnosis was the only matching trait available in this cohort. Individual measures of neuropathology were not comparable between cohorts, and cognitive assessments were not available for the Mayo cohort. A total of 257 metabolites of the 298 significant in the ROS/MAP cohort (across all 8 AD-related traits) were measured in the Mayo cohort. 30 of these 257 metabolites were significant in both datasets, i.e., with AD diagnosis in the Mayo cohort and with at least one of the eight AD-related traits in ROS/MAP (**Figure 3a**), all of which showed consistent effect directions.

**Figure 3:**
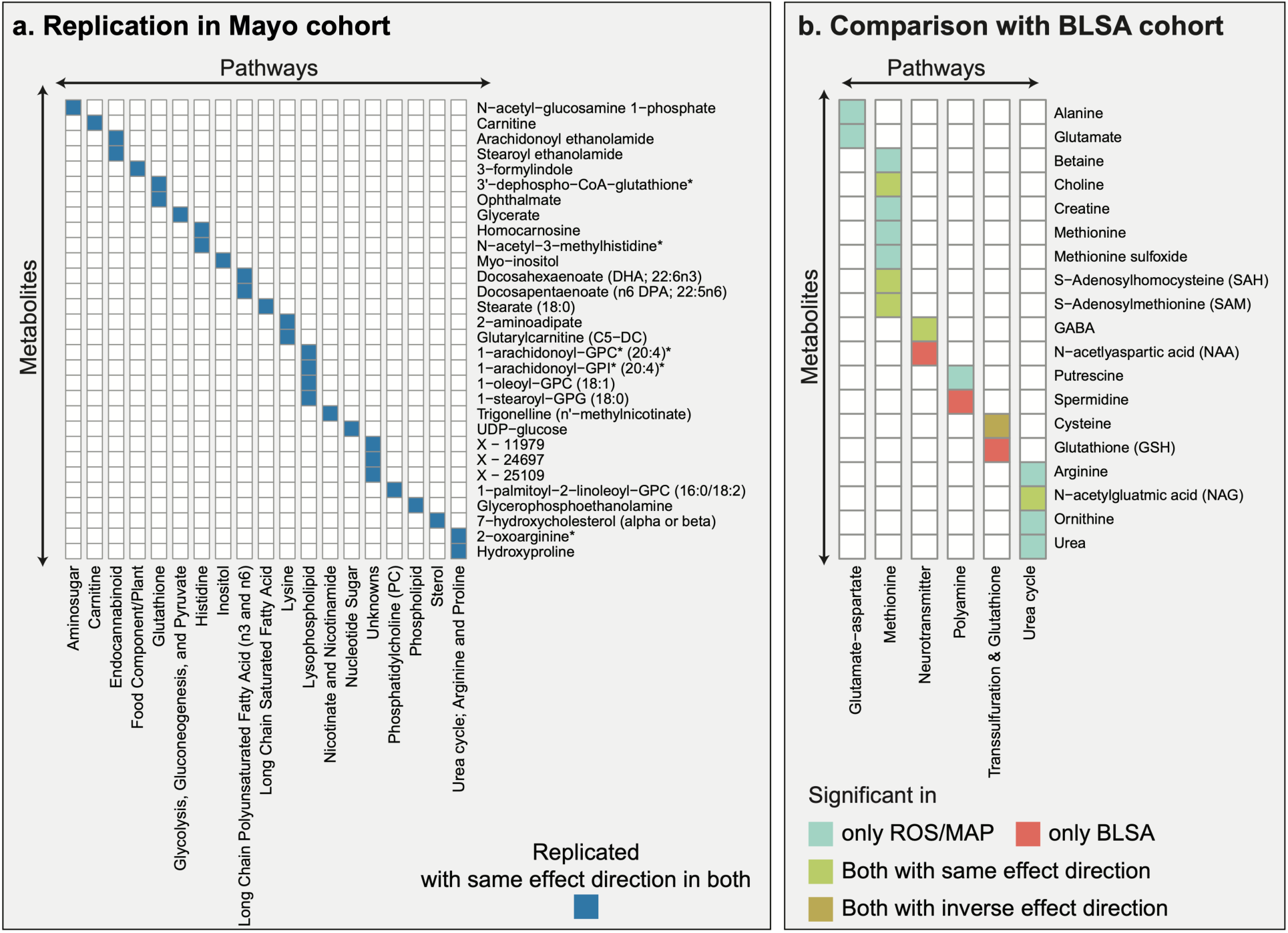
Overlap across independent cohorts and brain regions. **a**, The Mayo and ROS/MAP cohorts have 30 metabolic associations with consistent effect directions in common. **b**, The BLSA and ROS/MAP cohorts have 5 metabolic associations with consistent effect directions in common (green and brown). Cysteine showed inconclusive effect directions, with a positive association with AD in the BLSA cohort and a negative association with AD traits in the ROS/MAP cohort.

In the BLSA study (5), 43 samples from the inferior temporal gyrus (ITG) and middle frontal gyrus (MFG) brain regions were used for targeted metabolic profiling. The study identified 130 metabolites, of which the authors focused on 26, which were further categorized into different biochemical groups. In their analysis, 9 out of 26 metabolites were associated with AD diagnosis. All 26 metabolites were measured in our study, 17 of these 26 associated with AD-related traits, and 6 of these 17 were among the 9 metabolites associated in their study **(Figure 3b).** Of those 6 metabolites, 5 had the same effect directions, while cysteine was found to be positively associated with AD in the BLSA study and negatively associated with AD traits in ROS/MAP. Overall, 35 of the 298 associations identified in ROS/MAP were confirmed with consistent effect directions in either the Mayo or the BLSA cohort.

### 3.4 Metabolic alterations further characterize pathways previously implicated in AD

To examine the contribution of metabolic alterations in previously reported AD-related pathogenic processes, we selected four pathway groups for further exploration (**Figure 4**). Comprehensive functional annotations based on Metabolon’s sub-pathways are available in **Supplementary Figure 2**. For each pathway group discussed below, we included metabolites associated with at least one of the eight AD-related traits.

**Figure 4:**
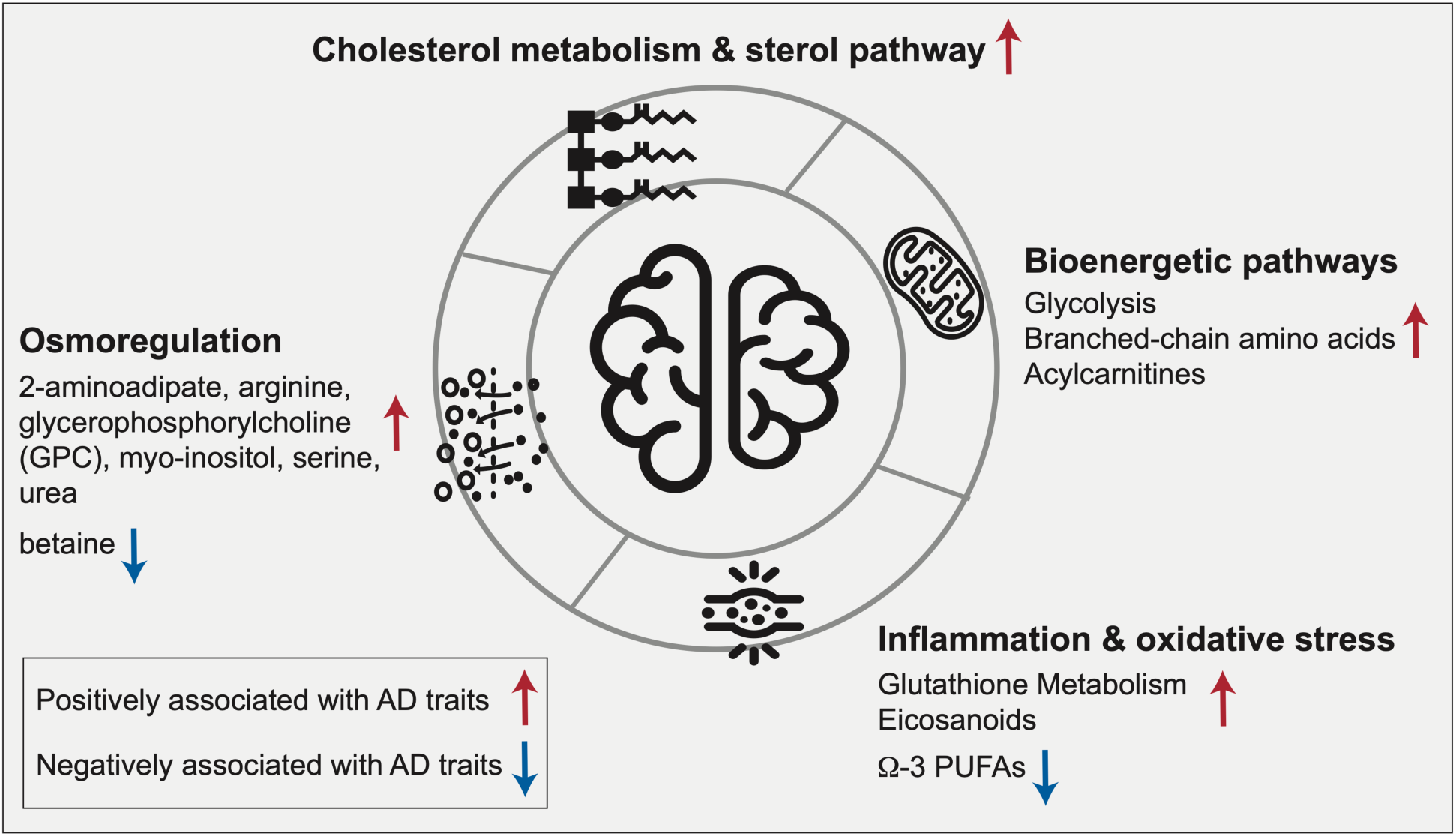
Pathway-level metabolic associations with AD-related traits. The highlighted biological processes have previously been implicated in AD. Our data provide a metabolic characterization of the AD-related alterations of these pathways in the brain: Cholesterol metabolism has an established connection to late-onset AD through *APOE*-ε4, the major genetic risk factor for the disease. Bioenergetic dysregulation is one of the earliest detectable changes in the central nervous system in AD and has also been described in the periphery. Inflammation and oxidative stress have been reported to synergistically affect AD pathogenesis. Osmoregulation affects various aspects of AD pathology, including protein folding, neural excitation, and autophagy.

#### 3.4.1 Bioenergetic pathways

Bioenergetic dysregulation is a hallmark of AD, which has been demonstrated using different technologies, from PET neuroimaging to deep molecular profiling such as metabolomics and proteomics studies (34–37). In our analysis, key metabolites from bioenergetic pathways, including glycolysis, branched-chain amino acid (BCAA) metabolism, and mitochondrial β-oxidation, were found to be significantly associated with AD. This included positive correlations with AD traits of the glycolytic metabolites glucose, glycerate, glucose 6-phosphate, and 1,5-anhydroglucitol; the BCAAs valine, isoleucine, and leucine, as well as their 1-carboxyethyl conjugates and degradation products β-hydroxyisovalerate, and 3-hydroxyisobutyrate; and the acylcarnitines isobutyrylcarnitine (C4), tiglyl carnitine (C5), 2-methylbutyrylcarnitine (C5), glutarylcarnitine (C5-DC), and 5-dodecenoylcarnitine (C12:1). High abundances of these metabolites resemble observations in blood metabolic profiles of individuals with type 2 diabetes and stand in contrast to blood-based studies in AD, which reported negative associations of, e.g., BCAA levels with AD (4, 38). Together, these results are in line with the hypothesis that AD might represent a “type 3” diabetes that selectively affects the brain (39).

#### 3.4.2 Cholesterol metabolism and sterol pathway

The strongest genetic risk for AD is exerted by variants of the *APOE* gene, a lipoprotein involved in cholesterol transport and metabolism (40–42). Our findings provide further evidence for the AD-associated significance of this pathway, with degradation products of cholesterol showing positive correlations with AD-related traits. These degradation products include 7α-hydroxy-3-oxo-4-cholestenoic acid (7-HOCA), which has been described as a CSF-based marker for blood-brain-barrier integrity (43); 4-cholesten-3-one, a product of cholesterol oxidation and isomerization through bacterial enzymes (44); and 7-hydroxycholesterol, a precursor for bile acids. Notably, cholesterol itself did not show any significant associations, indicating potential dysfunctional cholesterol clearance rather than a direct role of cholesterol in AD. This hypothesis is further supported by previous studies where we observed a significant increase of secondary bile acids in AD (39,45,46).

#### 3.4.3 Neuroinflammation and oxidative stress

Neuroinflammation is a central pathogenic feature of AD and is accompanied by the production of reactive oxygen species leading to oxidative stress (47). AD has been associated with both lipid mediators of inflammatory processes as well as immune response, including eicosanoids, and molecules involved in the antioxidant defense, such as glutathione (48–50). In line with these findings, we observed significant positive correlations of metabolites in the glutathione pathway with AD, indicating an upregulated antioxidant response. Significant metabolites included 4-hydroxy-nonenal-glutathione, a marker for detoxification of lipid peroxidation through glutathione S-transferases (GSTs) (51); cysteinylglycine disulfide, a degradation product of oxidized glutathione (49); and ophthalmate, an endogenous analog of hepatic glutathione (GSH) and a potential marker for GSH depletion (52). Moreover, pro-inflammatory eicosanoids showed positive associations with AD, including 15-oxoeicosatetraenoic acid (15-KETE), which has been linked to GST inhibition (53), and 12-hydroxy-heptadecatrienoic acid (12-HHTrE), overall providing further molecular evidence for active inflammatory processes in AD. In contrast, anti-inflammatory long-chain omega-3 polyunsaturated fatty acids (PUFAs), such as eicosapentaenoate (EPA) and docosahexaenoate (DHA) (54, 55), were negatively associated with AD.

#### 3.4.4 Osmoregulation

Osmolytes are a class of molecules that primarily sustain cell integrity (56). They have been suggested to play a neuroprotective role in AD by activating mTOR-independent autophagy signaling to inhibit the accumulation of aggregated proteins(57). Osmolytes also affect protein folding (58), and their therapeutic potential has been discussed in AD as well as other neurodegenerative proteinopathies (59). Moreover, osmolyte imbalances can impact neuronal hyperexcitation by influencing neurotransmitter uptake (56). In our analysis, we observed positive associations of several osmolytes with AD, including 2-aminoadipate, arginine, glycerophosphorylcholine (GPC), myo-inositol, serine, and urea, whereas betaine was negatively associated with the disease. As these observations are based on bulk tissue metabolomics, it remains unclear if these metabolites are deregulated within or outside of the cell. Nevertheless, the strong statistical significance underlying these associations suggests an important role of osmoregulation in AD which warrants further investigation.

### 3.5 Integration of complementary omics provides comprehensive view of biochemical cascade downstream of neurotransmitters

As a detailed showcase of the complex, biochemical interconnections in brain omics data, we selected a biochemical cascade downstream of the neurotransmitters glutamate and gamma aminobutyric acid (GABA). An elevated synaptic excitatory/inhibitory (E/I) ratio of these neurotransmitters has been linked to hyperexcitability and cognitive impairment observed in AD (60, 61). Furthermore, given GABA’s positive correlation with efficient working memory within the DLPFC region (62), it is of high significance to investigate GABA-related deregulation in this region.

We compiled biochemical steps of metabolites and enzymes downstream of glutamate using known reactions from the public database pathbank (63) (**Figure 5**). Notably, the cascade does not contain the routes from GABA to glutamate or from putrescine to GABA due to a lack of coverage of metabolites along those pathways.

**Figure 5:**
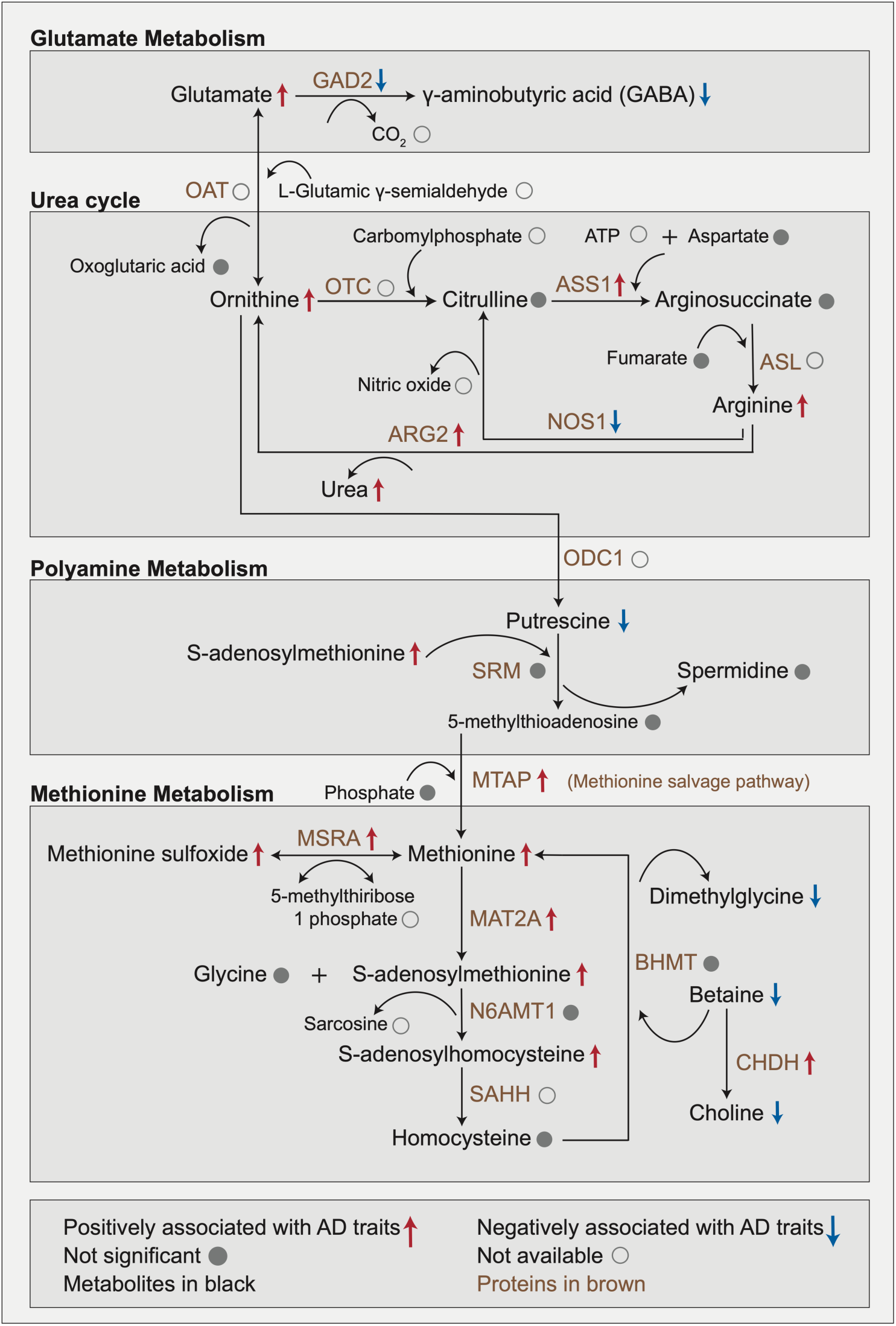
Metabolic changes downstream of the neurotransmitters glutamate/GABA. This multi-omics cascade starts with a biochemical process involving the conversion of glutamate into GABA within glutamate metabolism. Glutamate metabolism feeds into the urea cycle by conversion of glutamate to ornithine. Urea buildup to neurotoxic levels has been observed in postmortem brains of Huntington’s disease and has furthermore been linked to dementia. The urea cycle connects to polyamine metabolism via the conversion of ornithine into putrescine. Putrescine promotes the clearance of apoptotic cells via efferocytosis, a mechanism affected in AD. Polyamine metabolism connects to methionine metabolism through the methionine salvage pathway. Methionine acts as an antioxidant and is a precursor of s-adenosylmethionine, which is a key methyl donor in brain cells and involved in the synthesis of the neurotransmitters dopamine, norepinephrine, and serotonin via the folate cycle.

Based on proteomics profiles available for 262 matching brain samples, we performed a targeted association analysis of AD-related traits and proteins that are enzymatically involved in this pathway cascade (**Supplementary Table 10)**. Significant metabolic and proteomic associations with at least one of the eight AD-related traits were annotated on the respective molecules within the cascade.

#### 3.5.1 Glutamate metabolism

The pathway cascade starts with glutamate, which was positively associated with AD traits in our data. Excitatory glutamatergic synapses involving N-methyl D-aspartate receptors (NMDAR) have previously been targeted by memantine to treat severe AD (64). Glutamate is the precursor of the inhibitory neurotransmitter GABA, which we found to be negatively associated with AD.

Interestingly, protein abundance of glutamate decarboxylase (GAD2), which catalyzes the production of GABA from glutamate was also negatively associated with AD pathology in our data. This negative association provides a potential explanation for the imbalance between the two neurotransmitters.

#### 3.5.2 Urea cycle

Glutamate metabolism is directly connected to the urea cycle, in which ornithine, arginine, and urea were positively associated with AD. Urea buildup to neurotoxic levels has been observed in postmortem brains of Huntington’s disease and has furthermore been linked to dementia (65). The key enzyme in this pathway, arginase (ARG2) was positively associated with AD in our data. It catalyzes the conversion of arginine to ornithine with urea as a byproduct and has been previously linked to AD pathology due to its involvement in microglial activation and autophagy (66–68). Moreover, the reduction of urea levels through the inhibition of arginase (ARG2) has been suggested as a promising target in the context of AD (69).

#### 3.5.3 Polyamine metabolism

The urea cycle further feeds into the polyamine pathway, in which putrescine was negatively associated with AD, while spermidine and spermine were not significantly associated with AD. Putrescine promotes the clearance of apoptotic cells via efferocytosis (70), a mechanism affected in AD and other neurodegenerative diseases (71). Notably, some previous studies in human samples as well as mouse models have implicated putrescine in the context of AD (72–75).

#### 3.5.4 Methionine metabolism

In the final part of our pathway cascade, the enzyme S-methyl-5’-thioadenosine phosphorylase (MTAP) links the polyamine pathway to methionine metabolism, in which methionine, methionine sulfoxide, s-adenosylmethionine, and s-adenosylhomocysteine were positively associated with AD, with concordant changes in protein levels of respective enzymes MTAP, mitochondrial peptide methionine sulfoxide reductase (MSRA) and methionine adenosyltransferase (MAT2A). In a previous study, we have shown that higher levels of methionine in CSF were associated with AD (76). Methionine acts as an antioxidant by forming methionine sulfoxide and is also a precursor of s-adenosylmethionine (77), which is a key methyl donor in brain cells and involved in the synthesis of the neurotransmitters dopamine, norepinephrine, and serotonin via the folate cycle (78).

Overall, our analysis provides an integrated, multi-omics view of neurotransmitter-related changes known to play a role in the pathogenesis of AD.

### 3.6 Conditional analysis suggests tau pathology as a driver of metabolic changes in brain

According to the β-amyloid hypothesis of AD, β-amyloid is key to AD pathogenesis (79). It is considered to influence the accumulation of tangles of phosphorylated tau as well as tangle-driven pathogenesis (80). As a result, β-amyloid has been the focus of most therapeutic approaches (81–85). However, recent evidence suggests that tau tangles might be acting independent of β-amyloid (86). To identify metabolic signatures specific to β-amyloid and tau tangles we performed conditional analyses by adjusting for the respective other neuropathology (**Figure 6a**). In our standard association analysis, i.e., without accounting for β-amyloid load, 188 metabolites were associated with tau tangle load. 119 out of these 188 associations were still significant after accounting for β-amyloid load in the statistical model. While 34 metabolites were associated with β-amyloid load in the standard association analysis, only one remained significant after accounting for tau tangle load. Details of the standard and conditional analysis are available in **Supplementary Table 5** and **Supplementary Table 11**, respectively. Taken together, this analysis suggests that metabolic associations of tau tangles are largely independent of β-amyloid load, while metabolic associations of β-amyloid load are confounded by tau tangle load.

**Figure 6:**
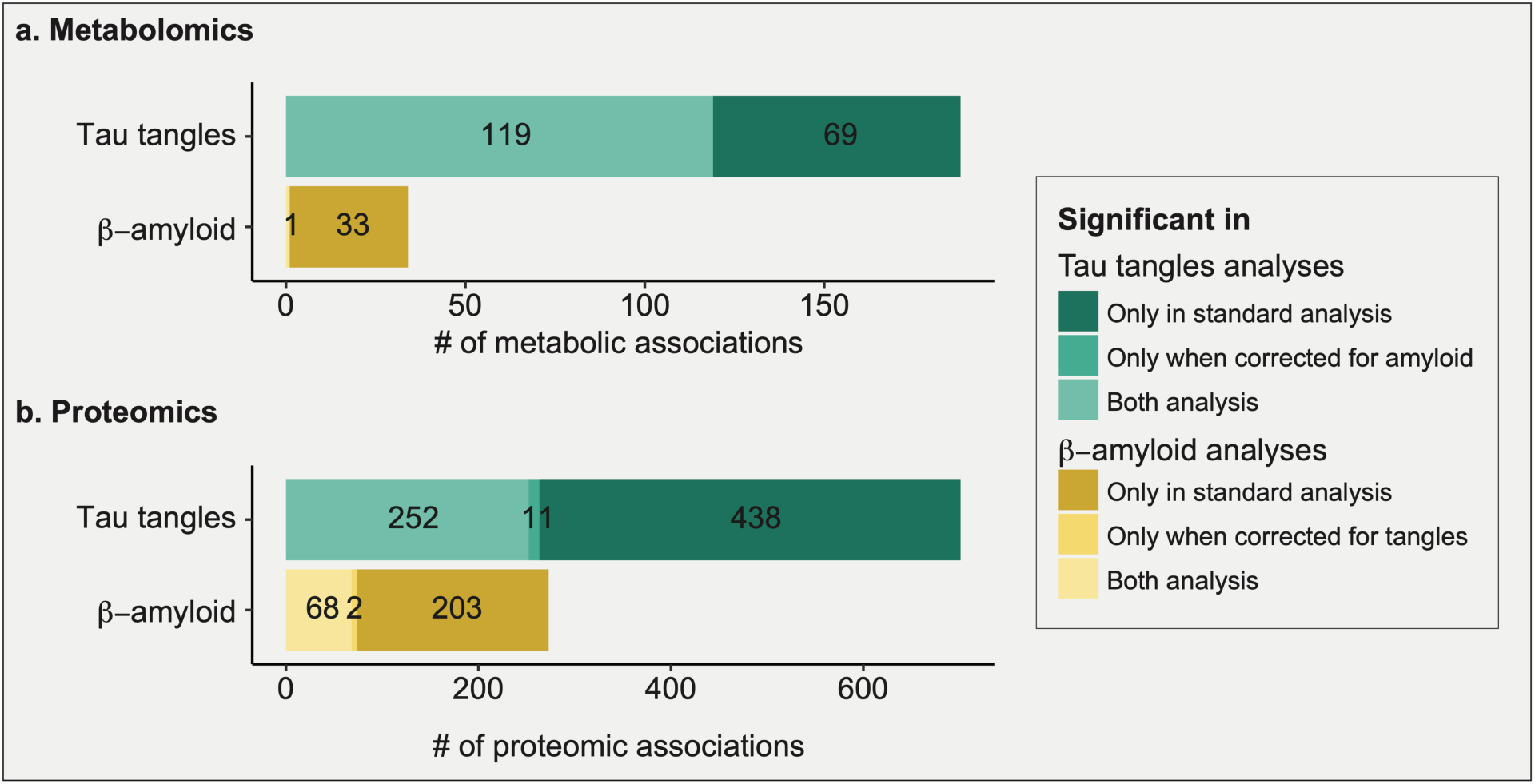
Comparison of standard and conditional analyses of β-amyloid and tau tangles. Tau tangle-associated signals appeared largely independent of β-amyloid signals, while β-amyloid signals were more strongly dependent on tau tangle signals. **a, b** Overlap of tangle- and β-amyloid-associated metabolites and proteins, respectively, with and without adjusting for the respective other neuropathology.

To corroborate this finding with another omics layer, we performed the same analysis on proteomics profiles. In the standard association analysis, 695 proteins associated with tau tangle load, i.e., without accounting for β-amyloid load. 252 out of these 695 were still associated with tau tangle load after accounting for β-amyloid load in our statistical model. While 265 proteins were associated with β-amyloid load in the standard association analysis, only 68 of these remained correlated after accounting for tau tangle load (**Figure 6b**). Details of the proteomics standard and conditional analysis are available in **Supplementary Table 12.**

Taken together, metabolic associations were more widespread for tau tangles and less dependent on β-amyloid load, which was partially confirmed by a similar trend in the proteomics data.

## 4 Discussion

In this work, we provide a global view of metabolic changes in brain related to Alzheimer’s disease. Our study is based on broad untargeted metabolomics profiles of 500 brain tissue samples from the DLPFC, covering 667 metabolites from various biochemical classes. We demonstrated that in cognitively normal individuals, age, sex, education, and BMI did not show major effects on brain metabolites. These limited associations of brain metabolites with demographics and socioeconomic status stand in contrast to the strong associations seen with blood metabolic profiles (28–30). Conversely, intake of medications had major effects on brain metabolome, as observed in blood metabolic profiles (4), highlighting the importance of accounting for the effects of pharmaceuticals. Of note, we observed significant modulation of metabolic associations through sex and *APOE-*ϕ.4 status, which is concordant with previous findings in blood-based metabolomics data (3).

### 4.1.1 Comparison with independent cohorts

In the subsequent association analysis, we found that 298 out of the 667 metabolites correlated with at least one of the eight investigated AD-related traits, covering cognition and several neuropathological parameters. We confirmed 30 of our associations using independent samples from the Mayo Clinic brain bank cohort. Additionally, 5 associations were confirmed using a study on the BLSA cohort (5). Two pathways, urea cycle, and glutathione metabolism, were associated with AD in all three cohorts. This overlap was observed despite the substantial differences in sample sizes, profiled brain regions, study designs, and clinical parameters. We thus conclude that the 35 metabolites and two pathways are high confidence AD-related metabolic signals in brain tissue, and the metabolic associations unique to our ROS/MAP study need further validation. Metabolic view of AD-associated pathways

Our study provided a metabolic view on various AD-related processes, including bioenergetic pathways, cholesterol metabolism, neuroinflammation, and osmoregulation. Out of those four, we will now specifically highlight our contribution towards the understanding of impaired bioenergetic metabolism and propose osmoregulation as a potentially important mechanism in AD pathogenesis. The bioenergetic alterations observed in our study endorse the existing evidence of impaired insulin signaling in AD (39). We speculate that this dysregulation (also referred to as type 3 diabetes) together with our finding that tau tangles are the major driver of metabolic alterations extends the emerging view that tau tangle is a key regulator of insulin signaling in the brain (87). Further, we observed numerous osmolytes being highly associated with AD, which suggests a potential functional link with the pathomechanisms of the disease. Osmolytes participate in multiple critical processes associated with neurogenerative diseases, including protein folding (88), autophagy (57), and hyperexcitation of neurons (56). While our observations on osmolytes might to some extent be confounded by, e.g., systematic differences in the hydration status of AD patients before death, to the best of our knowledge, these alterations within the central nervous system have so far not been studied in detail.

### 4.1.2 Biochemical changes downstream of glutamate/GABA neurotransmitters

We investigated detailed biochemical alterations downstream of glutamate and GABA. Imbalances of these neurotransmitters have previously been associated with hyperexcitability and cognitive impairment in AD (60, 61). In our study, the excitatory neurotransmitter glutamate was positively associated with AD, while the inhibitory neurotransmitter GABA was negatively associated with AD. These findings are in line with prior studies of AD etiology involving excess glutamate-mediated overstimulation (64) and emerging evidence that a decline in GABA levels contributes to synaptic dysfunction and excitatory/inhibitory imbalance (89). To investigate the downstream effects of this excitatory/inhibitory imbalance, we explored the metabolic and enzymatic changes in the biochemical cascade starting from the conversion of glutamate to GABA, connecting glutamate to urea cycle, polyamine metabolism, and methionine metabolism. Our study shows how the integration of metabolomics with proteomics provides a comprehensive overview of biochemical changes downstream of these neurotransmitters. Moreover, to the best of our knowledge, this is the first study reporting low levels of GABA being associated with cognitive decline within the DLPFC region. DLPFC is associated with working memory in individuals (62), which becomes impaired during AD pathogenesis (90). Thus, GABA levels within the DLPFC region have been of considerable interest to the AD community (90), which is corroborated by our results.

### 4.1.3 Interplay between proteinopathies and metabolic changes

Addressing the complex interplay of β-amyloid and tau tangles in AD, we performed a conditional statistical analysis. In our data, 97% of the β-amyloid-associated metabolites were dependent on tau tangle load, while only 36.7% of the tau tangle-associated metabolites were dependent on β-amyloid load. Our study thus provides preliminary evidence that the metabolic component of tau tangle-driven pathogenesis is independent of β-amyloid, which is in line with recent literature that proposes that tau accumulation might be independent of β-amyloid (86). Our finding may also suggest that metabolic changes in the brain are mostly later events in the pathologic cascade of AD (91) and closer temporally to tau pathology, neurodegeneration, and cognitive decline than to β-amyloid accumulation. Further supporting this, the largest number of associations in ROS/MAP was detected with cognitive decline, an event deemed to be at the later stages of the pathologic cascade of events in AD (92). Notably, our results might also have implications for potential pharmaceutical interventions for the disease. Specifically, it can be postulated that drugs targeting β-amyloid will only have limited impact on the metabolic component of AD.

### 4.1.4 Limitations

Despite the various novel insights into metabolic alterations in brain observed in AD, our study has several limitations. First, post-mortem studies are always cross-sectional in nature. The caveat of such studies is that they cannot be used to assess the causal direction of the identified associations. That is, an observed metabolic change in AD could be a factor directly contributing to disease development, or it could be a downstream effect of the pathological changes in brain. The true effect direction can only be determined in mechanistic follow-up studies or by genetic causality analysis such as Mendelian randomization (93), for which our study did not have the necessary statistical power. Second, diet and the gut microbiome are substantial confounders of the measured metabolites (94, 95), and examining the influence of these food-related factors will be a valuable and necessary addition in future studies. Third, the bulk tissue approach used in our study generates mixed metabolomic data from a variety of cell types and tissue compartments. Future work with specific cell populations will allow us to obtain a more precise picture of a particular mechanistic change accompanying AD and dementia. Fourth, postmortem tissue samples are prone to substantial biological and technical variation, as seen in the association of 307 out of 667 metabolites with postmortem interval (PMI), i.e., the time between death and sample preservation. Despite the statistical correction for PMI interval, degradation of certain metabolites until sample preservation is a factor that cannot be controlled in this type of study.

### 4.1.5 Conclusion and Outlook

Follow-up studies will be needed to build upon our findings, to complete the picture of dysregulated metabolism and pathological pathways in the Alzheimer’s disease brain. In particular, the wide availability of multi-omics datasets will provide a more holistic picture of the molecular changes associated with the disease (96–98). The integration of proteomics data into the glutamate/GABA pathway exploration in our study represents a pilot analysis in this direction; however, large-scale studies with an “ome-wide” integration of the (epi-)genome, transcriptome, proteome, and metabolome are required to further elucidate the mechanistic basis of AD pathogenesis and outline potential treatment options. To further enable these efforts, we published raw and processed metabolomics data through the AD Knowledge Portal, provide all analysis codes, and developed a reference catalog of hundreds of associations in an interactive web portal.

## Data availability

The data used in this paper can be obtained from two sources: (1) Metabolomics data for the ROS/MAP and Mayo cohorts, clinical data for the Mayo cohort, and proteomics data for the ROS/MAP cohort are available via the AD Knowledge Portal (https://adknowledgeportal.org). The AD Knowledge Portal is a platform for accessing data, analyses, and tools generated by the Accelerating Medicines Partnership (AMP-AD) Target Discovery Program and other National Institute on Aging (NIA)-supported programs to enable open-science practices and accelerate translational learning. The data, analyses, and tools are shared early in the research cycle without a publication embargo on secondary use. Data is available for general research use according to the following requirements for data access and data attribution (https://adknowledgeportal.org/DataAccess/Instructions). For access to content described in this manuscript see: http://doi.org/10.7303/syn26401311. (2) The full complement of clinical and demographic data for the ROS/MAP cohort are available via the Rush AD Center Resource Sharing Hub and can be requested at https://www.radc.rush.edu.

An interactive network view of AD associations from this study can be found at https://omicscience.org/apps/brainmwas/.

All R scripts to generate the tables and figures of this paper are available at https://github.com/krumsieklab/ad-brain-landscape.

## Acknowledgments

This work was done as part of the National Institute of Aging’s Accelerating Medicines Partnership for AD (AMP-AD) and was supported by NIH grants 1U19AG063744, 1R01AG069901-01A1, U01AG061357, P30AG10161, P30AG72975, R01AG15819, R01AG17917, U01AG46152, U01AG61356, RF1AG058942, RF1AG059093, and U01AG061359. The results published here are in whole or in part based on data obtained from the AD Knowledge Portal (https://adknowledgeportal.org).

The Religious Orders and the Rush Memory and Aging studies were supported by the National Institute on Aging grants P30AG10161, R01AG15819, R01AG17917, U01AG46152, and U01AG61356. The NIA also supported the Alzheimer Disease Metabolomics Consortium which is a part of the NIA’s national initiatives AMP-AD and M^2^OVE-AD (R01 AG046171, RF1 AG051550, and 3U01 AG061359-02S1). We thank the participants of ROS and MAP for their essential contributions and the gifts of their brains to these projects. All subjects gave informed consent.

The Mayo Clinic samples are part of the RNAseq study data led by Dr. Nilüfer Ertekin-Taner, Mayo Clinic, Jacksonville, FL as part of the multi-PI U01 AG046139 (MPIs Golde, Ertekin-Taner, Younkin, Price). Samples were provided from the following sources: The Mayo Clinic Brain Bank. Data collection was supported through funding by NIA grants P50 AG016574, R01 AG032990, U01 AG046139, R01 AG018023, U01 AG006576, U01 AG006786, R01 AG025711, R01 AG017216, R01 AG003949, NINDS grant R01 NS080820, CurePSP Foundation, and support from Mayo Foundation.

RB thanks her colleagues from the Krumsiek lab for fruitful discussions and support in this work.

## Author contributions

RB, MArnold, GK, JK designed the computational and statistical methods, performed the analysis, interpreted the results, and drafted the manuscript. MAW created the interactive online supplement. DAB is PI of the ROS/MAP study and provided samples as well as phenotypic data. NET, XW, MAllen provided samples and phenotypic data on the Mayo clinic cohort and contributed to the analysis of those samples. CB curated and managed phenotypic data. AIL, NTS provided the preprocessed proteomics data for matching ROS/MAP samples and contributed to the analysis of those samples. All authors read and reviewed the manuscript. RK-D acquired funding and is the overall PI of the Alzheimer’s disease metabolomics consortium.

