## Supplementary Figure 1 for "The landscape of metabolic brain alterations in Alzheimer’s disease"

**Supplementary Figure 1: Distribution and overlap of metabolic associations across the AD-related traits.** Vertical bars in upper half indicate the size of the set represented by the dots in the lower half. Horizontal bars in the lower left quadrant indicate number of associations per trait. Single dots in the lower panel represent unique hits to that trait.

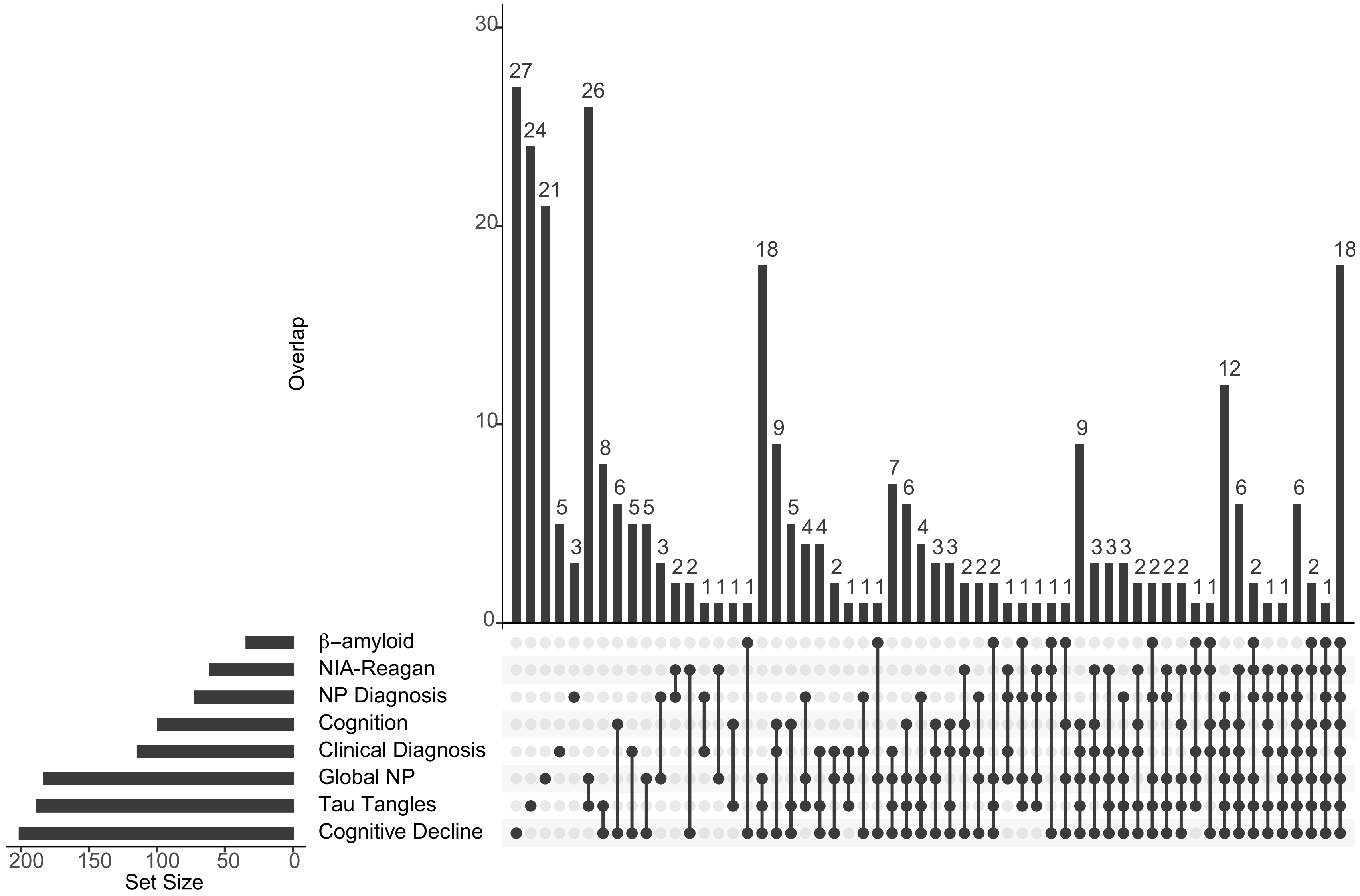
