## Supplementary Figure 2 for "The landscape of metabolic brain alterations in Alzheimer’s disease"

### Supplementary Figure 2:
### Distribution of metabolic associations across sub-pathways

### of significant metabolites: 0 – 19 – 38

| Super-pathway | β–amyloid | Tau tangles | Global NP | NIA-Reagan | NP Diagnosis | Clinical Diagnosis | Cognitive Decline | Cognition | Total | Sub-Pathways |
|---|---|---|---|---|---|---|---|---|---|---|
| Amino Acid | 2 | 2 | 3 | 1 | 1 | 2 | 2 | 3 | 3 | Alanine and Aspartate Metabolism |
| Amino Acid | 0 | 1 | 2 | 1 | 0 | 1 | 1 | 3 | 3 | Creatine Metabolism |
| Amino Acid | 3 | 5 | 7 | 3 | 3 | 6 | 7 | 9 | 9 | Glutamate Metabolism |
| Amino Acid | 0 | 2 | 0 | 0 | 0 | 1 | 1 | 3 | 4 | Glutathione Metabolism |
| Amino Acid | 2 | 6 | 6 | 2 | 3 | 4 | 5 | 6 | 8 | Glycine, Serine and Threonine Metabolism |
| Amino Acid | 1 | 2 | 6 | 0 | 4 | 2 | 1 | 4 | 7 | Histidine Metabolism |
| Amino Acid | 1 | 8 | 9 | 2 | 1 | 5 | 3 | 7 | 11 | Leucine, Isoleucine and Valine Metabolism |
| Amino Acid | 1 | 5 | 5 | 1 | 1 | 4 | 4 | 4 | 6 | Lysine Metabolism |
| Amino Acid | 1 | 4 | 2 | 2 | 1 | 4 | 2 | 3 | 8 | Methionine, Cysteine, SAM and Taurine Metabolism |
| Amino Acid | 0 | 2 | 2 | 2 | 1 | 2 | 2 | 2 | 2 | Phenylalanine Metabolism |
| Amino Acid | 1 | 3 | 3 | 1 | 2 | 4 | 4 | 4 | 4 | Polyamine Metabolism |
| Amino Acid | 0 | 2 | 2 | 2 | 0 | 1 | 1 | 2 | 2 | Tryptophan Metabolism |
| Amino Acid | 1 | 2 | 3 | 3 | 1 | 2 | 1 | 2 | 3 | Tyrosine Metabolism |
| Amino Acid | 1 | 5 | 5 | 1 | 2 | 4 | 2 | 5 | 8 | Urea cycle; Arginine and Proline Metabolism |
| Carbohydrate | 0 | 1 | 2 | 0 | 1 | 1 | 1 | 3 | 4 | Aminosugar Metabolism |
| Carbohydrate | 0 | 0 | 1 | 0 | 0 | 0 | 0 | 0 | 1 | Disaccharides and Oligosaccharides |
| Carbohydrate | 0 | 1 | 0 | 0 | 0 | 1 | 0 | 2 | 2 | Fructose, Mannose and Galactose Metabolism |
| Carbohydrate | 0 | 2 | 0 | 0 | 0 | 0 | 0 | 0 | 2 | Glycogen Metabolism |
| Carbohydrate | 0 | 3 | 1 | 0 | 0 | 2 | 2 | 3 | 5 | Glycolysis, Gluconeogenesis, and Pyruvate Metabolism |
| Carbohydrate | 0 | 1 | 1 | 1 | 1 | 1 | 1 | 1 | 1 | Nucleotide Sugar |
| Carbohydrate | 0 | 1 | 1 | 0 | 0 | 1 | 0 | 3 | 3 | Pentose Metabolism |
| Carbohydrate | 0 | 0 | 1 | 1 | 1 | 1 | 0 | 0 | 1 | Pentose Phosphate Pathway |
| Cofactors and Vitamins | 1 | 2 | 2 | 0 | 2 | 1 | 1 | 2 | 2 | Ascorbate and Aldarate Metabolism |
| Cofactors and Vitamins | 0 | 1 | 1 | 0 | 0 | 0 | 0 | 1 | 1 | Hemoglobin and Porphyrin Metabolism |
| Cofactors and Vitamins | 0 | 5 | 3 | 0 | 2 | 4 | 3 | 4 | 5 | Nicotinate and Nicotinamide Metabolism |
| Cofactors and Vitamins | 0 | 1 | 1 | 0 | 1 | 0 | 0 | 1 | 1 | Riboflavin Metabolism |
| Cofactors and Vitamins | 0 | 0 | 0 | 0 | 0 | 1 | 1 | 1 | 1 | Tocopherol Metabolism |
| Cofactors and Vitamins | 0 | 0 | 0 | 0 | 0 | 1 | 1 | 1 | 1 | Vitamin A Metabolism |
| Cofactors and Vitamins | 0 | 0 | 1 | 0 | 0 | 0 | 0 | 0 | 1 | Vitamin B6 Metabolism |
| Energy | 0 | 1 | 2 | 1 | 0 | 1 | 1 | 1 | 2 | TCA Cycle |
| Lipid | 1 | 2 | 2 | 2 | 2 | 1 | 1 | 2 | 2 | Carnitine Metabolism |
| Lipid | 0 | 5 | 2 | 0 | 0 | 0 | 0 | 1 | 6 | Ceramides |
| Lipid | 0 | 0 | 0 | 0 | 1 | 0 | 0 | 0 | 1 | Corticosteroids |
| Lipid | 0 | 1 | 0 | 0 | 0 | 0 | 0 | 1 | 1 | Diacylglycerol |
| Lipid | 0 | 2 | 2 | 0 | 0 | 0 | 0 | 0 | 2 | Dihydroceramides |
| Lipid | 1 | 2 | 2 | 1 | 1 | 0 | 0 | 1 | 2 | Dihydrosphingomyelins |
| Lipid | 0 | 0 | 0 | 1 | 0 | 1 | 1 | 2 | 2 | Eicosanoid |
| Lipid | 2 | 2 | 2 | 1 | 1 | 0 | 1 | 3 | 3 | Endocannabinoid |
| Lipid | 0 | 0 | 0 | 0 | 0 | 0 | 0 | 1 | 1 | Fatty Acid Metabolism (Acyl Carnitine, Long Chain Saturated) |
| Lipid | 0 | 0 | 0 | 0 | 0 | 0 | 0 | 2 | 2 | Fatty Acid Metabolism (Acyl Carnitine, Monounsaturated) |
| Lipid | 0 | 0 | 0 | 0 | 0 | 0 | 1 | 1 | 2 | Fatty Acid Metabolism (Acyl Carnitine, Polyunsaturated) |
| Lipid | 0 | 0 | 0 | 0 | 0 | 0 | 0 | 1 | 1 | Fatty Acid, Amino |
| Lipid | 0 | 2 | 2 | 0 | 0 | 0 | 1 | 1 | 2 | Fatty Acid, Branched |
| Lipid | 0 | 2 | 2 | 2 | 1 | 2 | 2 | 3 | 3 | Fatty Acid, Dicarboxylate |
| Lipid | 0 | 1 | 2 | 0 | 1 | 1 | 0 | 1 | 2 | Fatty Acid, Dihydroxy |
| Lipid | 0 | 1 | 0 | 1 | 0 | 1 | 1 | 1 | 1 | Fatty Acid, Monohydroxy |
| Lipid | 0 | 1 | 1 | 0 | 0 | 0 | 0 | 0 | 1 | Glycerolipid Metabolism |
| Lipid | 0 | 2 | 1 | 0 | 0 | 0 | 0 | 0 | 3 | Hexosylceramides (HCER) |
| Lipid | 1 | 2 | 2 | 2 | 1 | 2 | 2 | 3 | 3 | Inositol Metabolism |
| Lipid | 0 | 1 | 0 | 0 | 0 | 0 | 1 | 1 | 1 | Lactosylceramides (LCER) |
| Lipid | 0 | 5 | 3 | 0 | 0 | 0 | 1 | 2 | 5 | Long Chain Polyunsaturated Fatty Acid (n3 and n6) |
| Lipid | 0 | 4 | 3 | 0 | 0 | 0 | 0 | 1 | 4 | Long Chain Saturated Fatty Acid |
| Lipid | 0 | 10 | 8 | 0 | 0 | 1 | 1 | 5 | 12 | Lysophospholipid |
| Lipid | 0 | 1 | 0 | 0 | 0 | 0 | 0 | 0 | 1 | Lysoplasmalogen |
| Lipid | 0 | 2 | 1 | 0 | 0 | 1 | 0 | 2 | 2 | Medium Chain Fatty Acid |
| Lipid | 0 | 9 | 12 | 2 | 4 | 5 | 2 | 7 | 15 | Phosphatidylcholine (PC) |
| Lipid | 2 | 7 | 8 | 2 | 8 | 6 | 3 | 7 | 10 | Phosphatidylethanolamine (PE) |
| Lipid | 0 | 2 | 2 | 0 | 1 | 0 | 0 | 1 | 2 | Phosphatidylglycerol (PG) |
| Lipid | 1 | 1 | 2 | 1 | 1 | 1 | 0 | 2 | 3 | Phosphatidylinositol (PI) |
| Lipid | 2 | 5 | 5 | 2 | 3 | 4 | 3 | 5 | 7 | Phospholipid Metabolism |
| Lipid | 0 | 1 | 1 | 1 | 1 | 1 | 0 | 1 | 1 | Plasmalogen |
| Lipid | 0 | 1 | 1 | 0 | 0 | 0 | 0 | 1 | 1 | Sphingolipid Synthesis |
| Lipid | 2 | 4 | 3 | 0 | 2 | 0 | 0 | 3 | 5 | Sphingomyelins |
| Lipid | 0 | 1 | 1 | 0 | 0 | 0 | 0 | 1 | 1 | Sphingosines |
| Lipid | 0 | 2 | 1 | 3 | 2 | 2 | 2 | 3 | 3 | Sterol |
| Nucleotide | 0 | 0 | 1 | 0 | 0 | 0 | 0 | 0 | 1 | Purine and Pyrimidine Metabolism |
| Nucleotide | 0 | 2 | 3 | 0 | 1 | 1 | 0 | 2 | 4 | Purine Metabolism, (Hypo)Xanthine/Inosine containing |
| Nucleotide | 1 | 2 | 1 | 0 | 0 | 0 | 1 | 3 | 3 | Purine Metabolism, Adenine containing |
| Nucleotide | 0 | 2 | 3 | 0 | 0 | 0 | 1 | 3 | 3 | Purine Metabolism, Guanine containing |
| Nucleotide | 0 | 0 | 1 | 0 | 0 | 1 | 0 | 1 | 2 | Pyrimidine Metabolism, Cytidine containing |
| Nucleotide | 0 | 1 | 1 | 1 | 0 | 0 | 1 | 1 | 1 | Pyrimidine Metabolism, Thymine containing |
| Nucleotide | 0 | 2 | 2 | 0 | 0 | 0 | 1 | 3 | 5 | Pyrimidine Metabolism, Uracil containing |
| Peptide | 0 | 3 | 2 | 0 | 0 | 0 | 0 | 2 | 3 | Dipeptide |
| Peptide | 0 | 2 | 1 | 0 | 0 | 2 | 1 | 1 | 3 | Gamma−glutamyl Amino Acid |
| Unknowns | 6 | 21 | 22 | 14 | 12 | 20 | 21 | 31 | 38 | Unknowns |
| Xenobiotics | 0 | 0 | 0 | 0 | 0 | 0 | 1 | 1 | 1 | Bacterial/Fungal |
| Xenobiotics | 0 | 1 | 0 | 0 | 0 | 0 | 0 | 2 | 2 | Benzoate Metabolism |
| Xenobiotics | 0 | 2 | 2 | 0 | 0 | 0 | 0 | 1 | 3 | Chemical |
| Xenobiotics | 0 | 1 | 1 | 1 | 1 | 4 | 2 | 4 | 5 | Food Component/Plant |

β–amyloid, Tau tangles, Global NP, NIA-Reagan, NP Diagnosis, Clinical Diagnosis, Cognition, Cognitive Decline, Total

**AD traits**

Sub-Pathways

Super-pathways: Amino Acid, Carbohydrate, Cofactors and Vitamins, Energy, Lipid, Nucleotide, Peptide, Unknowns, Xenobiotics
